# Maternal immune activation dysregulation of the fetal brain transcriptome and relevance to the pathophysiology of autism spectrum disorder

**DOI:** 10.1101/064683

**Authors:** Michael V. Lombardo, Hyang Mi Moon, Jennifer Su, Theo D. Palmer, Eric Courchesne, Tiziano Pramparo

## Abstract

Maternal immune activation (MIA) via infection during pregnancy is known to increase risk for autism spectrum disorder (ASD). However, it is unclear how MIA disrupts fetal brain gene expression in ways that may explain this increased risk. Here we examine how MIA dysregulates fetal brain gene expression near the end of the first trimester of human gestation in ways relevant to ASD-associated pathophysiology. MIA downregulates expression of ASD-associated genes, with the largest enrichments in genes known to harbor rare highly penetrant mutations. MIA also downregulates expression of many genes also known to be persistently downregulated in ASD cortex later in life and which are canonically known for roles in affecting prenatally-late developmental processes at the synapse. Transcriptional and translational programs that are downstream targets of highly ASD-penetrant *FMR1* and *CHD8* genes are also heavily affected by MIA. MIA strongly upregulates expression of a large number of genes involved in translation initiation, cell cycle, DNA damage, and proteolysis processes that affect multiple key neural developmental functions. Upregulation of translation initiation is common to and preserved in gene network structure with the ASD cortical transcriptome throughout life and has downstream impact on cell cycle processes. The cap-dependent translation initiation gene, *EIF4E*, is one of the most MIA-dysregulated of all ASD-associated genes and targeted network analyses demonstrate prominent MIA-induced transcriptional dysregulation of *mTOR* and *EIF4E*-dependent signaling. This dysregulation of translation initiation via alteration of the *Tsc2-mTor*-*Eif4e*-axis was further validated across MIA rodent models. MIA may confer increased risk for ASD by dysregulating key aspects of fetal brain gene expression that are highly relevant to pathophysiology affecting ASD.

---

Multiple etiological pathways contribute to increased risk for autism spectrum disorder (ASD). For example, there are now a handful of identified rare de novo variants with high penetrance for ASD^1-4^, with theoretically many others that have yet to be discovered^5^. Interestingly, such rare high-confidence mutations tend to be significantly enriched in genes involved in synaptic functions, transcriptional regulation, and chromatin remodeling functions and/or are downstream targets of the fragile X syndrome protein (FMRP) complex^1,3^. In contrast, common variants may also significantly contribute to a large proportion (up to 60%) of genetic liability for ASD^6,7^, suggesting that hundreds of genes, individually associated with a small risk, may underlie ASD etiology via a much larger collective effect that acts at the network level either alone or in combination with environmental factors. Supporting this model, evidence from twin studies suggest that while heritability is quite high^8^, there is also a substantial environmental component for ASD susceptibility^9^. Recent evidence^10-19^ has also catalyzed the concept that genetic and non-genetic factors and their interaction, may act at very early periods of fetal brain development and potentially alter protein or gene expression regulation leading to potentially shared pathways for complex ASD-related phenotypes. Thus, much can be learned about the biological processes and molecular mechanisms involved in ASD by modeling environmental risk factors and studying their effects on functional genomics during early developmental stages of fetal brain development.

One environmental factor known to alter early fetal brain development and increase the risk for ASD is maternal infection during pregnancy^16-18, 20-23^. Prenatal maternal infection on fetal brain development can be studied with maternal immune activation (MIA) animal models^24,25^. MIA induces maternal cytokine signaling that passes through the placenta to affect fetal brain development^26^ and blocking key pathways prevents MIA-induced neural and behavioral abnormalities in ASD model systems^27^. The consequences of MIA include behavioral deficits of broad relevance to ASD^28-30^ as well as numerous ASD-relevant influences on the developing brain^31^. These influences include upregulation of cell cycle gene expression^26^ and shortening of cell cycle as seen in ASD^32^, over-production of neurons^33^ analogous to some cases of ASD^13^, increased cortical thickness^33^, increased brain size^34^ as seen in many ASD toddlers^35,36^, altered expression of genes involved in neuronal migration^26^, cortical layering defects^37^ including focal patches of disorganized cortex^27^ analogous to reports in some ASD cases^10^, microglia abnormalities and enhanced microglia priming^34, 38^ as seen in ASD^39-41^, alteration of GABAergic signaling^42^, cerebellar vermis defects^43^, and defects of prefrontal dendritic morphology^44^.

Despite the numerous links between MIA and ASD pathology, several key questions remain with regards to how MIA affects the developing fetal brain at a genomic and epigenomic levels and how such influence maps onto known genetic risk mechanisms associated with ASD. For example, does MIA exert its influence via genes associated with ASD and if so, which classes of genetic variants are most highly affected? Can MIA induce transcriptomic pathology in the fetal brain that shares similarities with cortical transcriptome dysregulation that is present in children and adults with ASD^45-47^? What functional genomic pathologies are present in the MIA-induced fetal brain that are not present in older children and adults with ASD? Are there specific mechanistic pathways that MIA dysregulates that are highly relevant for ASD? A better understanding of these key mechanistic links can help to further understand how MIA may confer risk for later development of ASD. By better understanding these mechanistic links between MIA and ASD, this work may ultimately help lead towards development of potential therapeutic targets for specific environmental risk factors that may be more amenable to prevention and/or treatment later in life^48,49^ than genetic etiologies. Furthermore, if MIA alters expression in pathways shared with those in Fragile X Syndrome for which advances in drug development are in progress, then drugs that successfully target those pathways in Fragile X could potentially be re-purposed.

In this work, we leverage bioinformatic and statistical approaches on available MIA gene expression data to investigate several key hypotheses about how MIA may dysregulate the fetal brain transcriptome in ways relevant to ASD. We first test the two hypotheses that MIA-induced effects may directly downregulate the expression of genes known to be associated with ASD and may indirectly alter protein targets downstream from two master regulatory genes of high penetrance for ASD (i.e. *FMR1* and *CHD8*). We then test the hypothesis that MIA dysregulates the fetal brain transcriptome in ways that are similar to cortical transcriptome dysregulation observed children and adults with ASD. We also heavily focus on how similarities in atypical biological systems in ASD and MIA can manifest in key pathways that are critically important for ASD and also reveal which early MIA-induced functional genomic pathologies are not detectable in the mature ASD brain. Finally, we independently induce MIA in mice to validate gene expression alterations in one prominent molecular pathway critical for protein translation processes during early fetal brain development and relevant to ASD pathophysiology.

## Results

We evaluated MIA-induced differential expression (DE) in a rat dataset from Oskvig et al.,^26^ measured at 4 hours post-lipopolysaccharide (LPS) injection on gestational day 15. This manipulation in rat corresponds roughly to post-conception day 68 of human prenatal cortical development, which is analogous to the end of the first trimester of pregnancy^50^ (Supplementary Fig 1). Generally consistent with analyses in Oskvig et al, here we found massive MIA transcriptome dysregulation of 4959 downregulated genes (5398 probes) and 4033 upregulated genes (4462 probes) (see Supp Table 1 for gene lists).

To describe processes enriched in such MIA DE gene sets we used MetaCore GeneGO for pathway analysis. MIA-downregulated genes displayed enriched functions relevant to both early cortical development - such as WNT /Hedgehog signaling and neurogenesis - and, later cortical development such as axonal guidance and synaptogenesis (Fig 1A). In contrast, MIA-upregulated genes displayed predominant enrichment in processes that can play key roles in neurogenesis and early brain development, such as translation, cell cycle, DNA damage, and proteolysis processes (Fig 2A) (see Supp Table 2 for full list of enrichments). This MIA-induced overexpression of early processes affecting protein synthesis, cell number, DNA integrity, and cell fate specification are in line with functions that would be expected to be normally active during late first trimester of human brain development and are consistent with some hypotheses of the early neural abnormalities in ASD such as dysregulated neurogenesis^10, 13, 31, 32, 51^.

**Figure 1:**
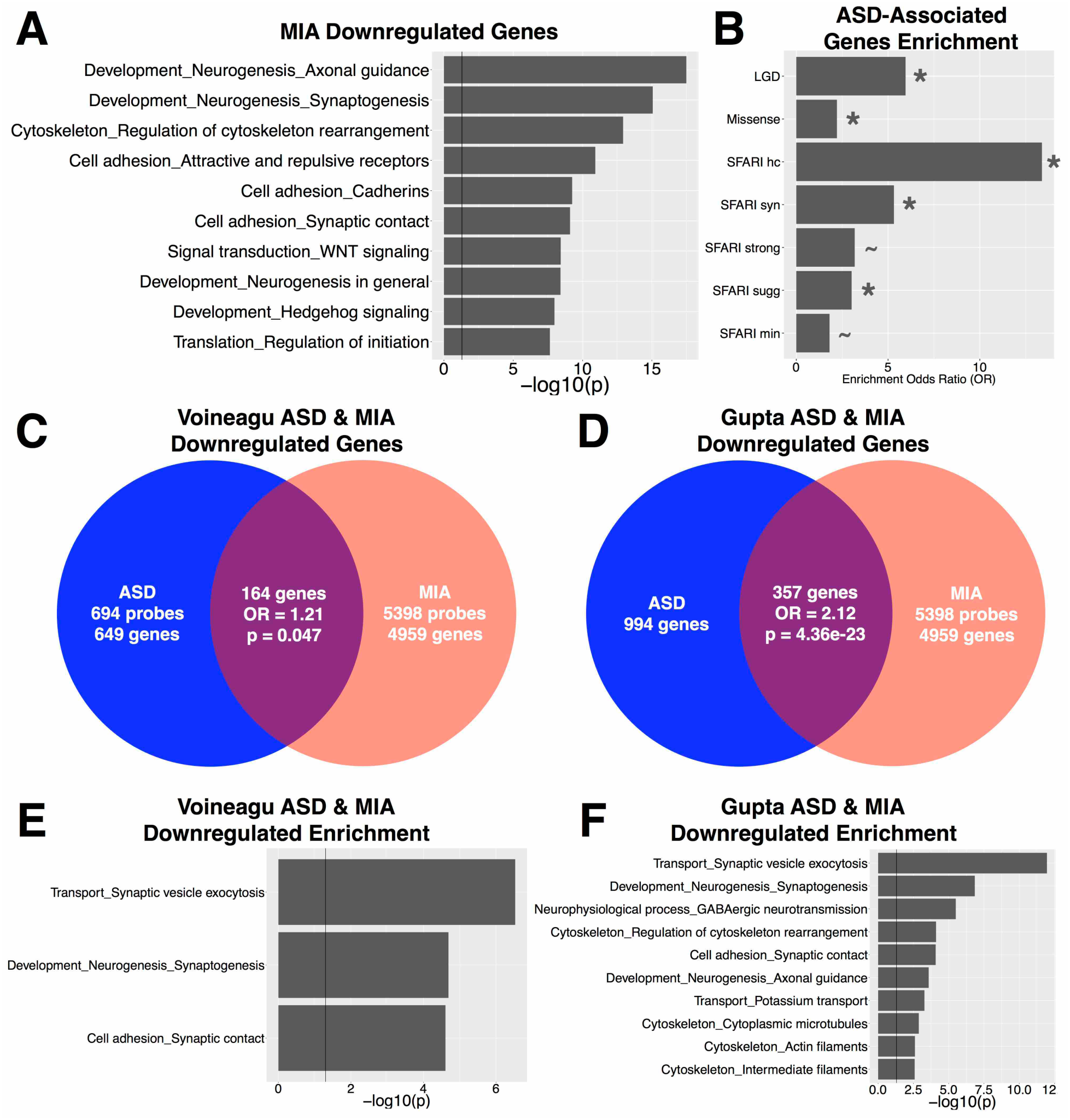
Enrichment of MIA-downregulated genes with classes of ASD-associated genes and ASD cortical transcriptome downregulated genes. This figure describes MIA-downregulated genes and their enrichment within different classes of ASD-associated genes and genes that downregulated in the ASD cortical transcriptome. Panel A shows process level enrichments for all MIA-downregulated genes. Panel B shows enrichment odds ratios for different classes of ASD-associated genes (the * indicates enrichment passing FDR q<0.05, while the ∼ indicates enrichment passing FDR q<0.1). Panels CD show enrichment between downregulated genes in MIA and ASD cortical transcriptome datasets (panel C for the Voineagu dataset, panel D for the Gupta dataset). Panels E-F show process level enrichments for the common downregulated genes between MIA and ASD (panel E for the Voineagu dataset, panel F for the Gupta dataset).

**Figure 2:**
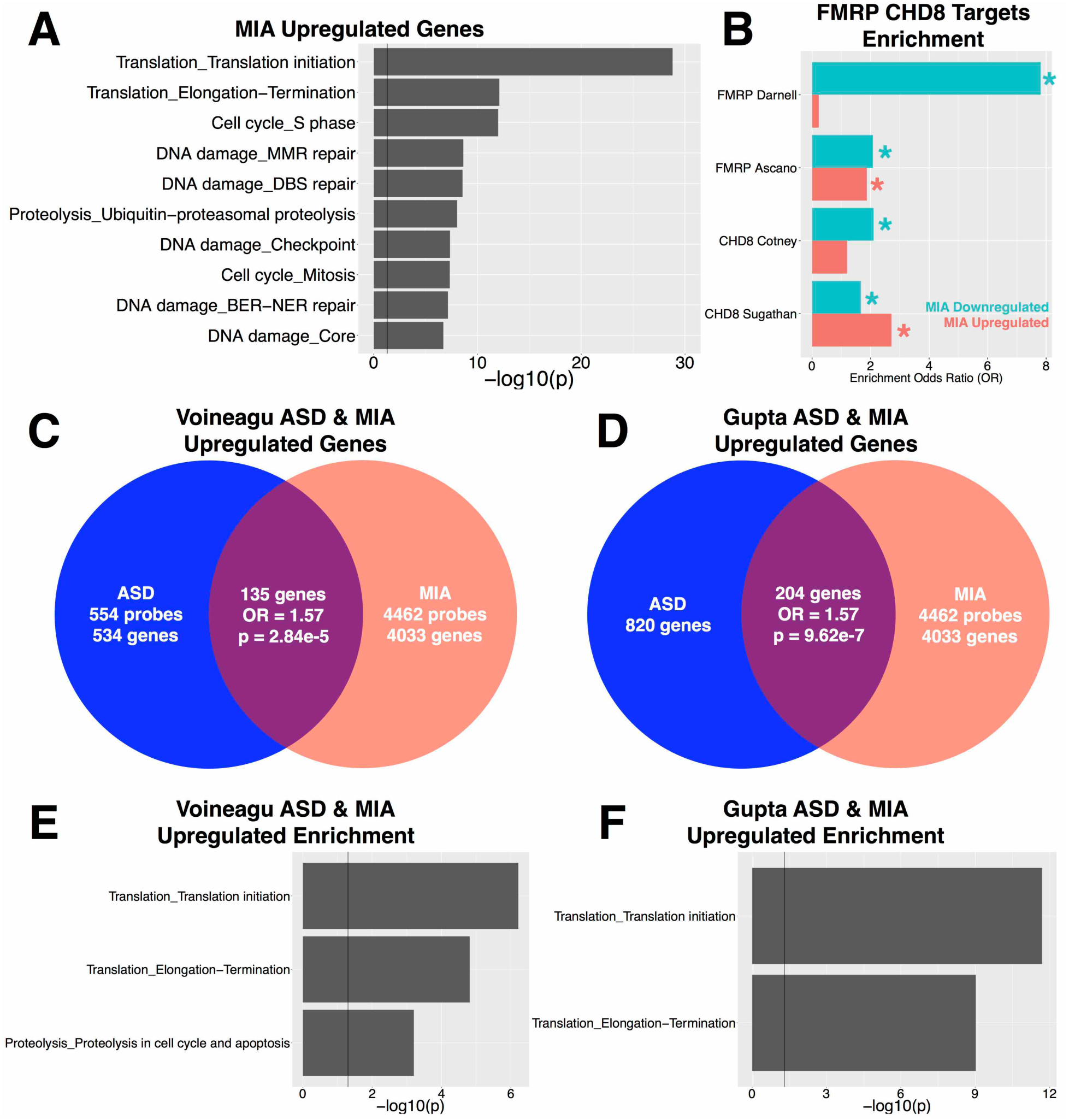
MIA effects on translation and transcriptional mechanisms. This figure shows results supporting the idea that MIA dysregulates processes involved in translation and transcription. Panel A shows process level enrichments for MIA-upregulated genes. Panel B shows enrichment odds ratios for MIA-downregulated or upregulated gene sets with downstream FMRP and CHD8 targets (* indicates enrichment test passing FDR q<0.05 threshold). Panels C-D show enrichment between upregulated genes in MIA and ASD cortical transcriptome datasets (panel C for the Voineagu dataset, panel D for the Gupta dataset). Panels E-F show process level enrichments for the common upregulated genes between MIA and ASD (panel E for the Voineagu dataset, panel F for the Gupta dataset).

### Classes of ASD-associated genes are enriched with MIA-downregulated genes

We next examined our first hypothesis that MIA-downregulated genes are substantially enriched in various classes of ASD-associated genes. Our first test of this hypothesis examined a class of genes identified by recent whole-exome sequencing studies as “likely gene-disrupting” variants (LGD) (i.e. splice-site, nonsense, or frameshift variants) (see Supp Table 1 for gene lists)^1-4^. Remarkably, 57% (20/35) of LGD genes were present in the MIA downregulated gene set, amounting to a substantial enrichment (OR = 5.95, p = 9.44e-6). When considering known ASD-associated missense variants^3^, we also found substantial enrichment (33%, 48/145, OR = 2.21, p = 0.0021) (Fig 1B). We then further considered ASD-associated gene classes separated by the expert manually-curated categories in the SFARI Gene database (http://gene.sfari.org/^52, 53^) (see Supp Table 1 for gene lists). Here we also found that the MIA-downregulated genes are substantially enriched in several categories, with a gradient in enrichment that follows the strength of evidence implied by each category. The strongest enrichments by enrichment odds ratio were within the SFARI High Confidence category (75%, 12/16, OR = 13.38, p = 1.18e-5), followed by Syndromic (54%, 25/46, OR = 5.32, p = 2.57e-6), Strong (41%, 10/24, OR = 3.18, p = 0.0277), and Suggestive gene categories (40%, 27/67, OR = 3.01, p = 7.91e-4) (Fig 1B). These findings suggest that MIA may increase risk for ASD via downregulating at a very early stage of brain development the expression of many of the same genes that are known to be highly penetrant for ASD.

### MIA dysregulates downstream targets of FMR1 and CHD8

The evidence that MIA downregulates expression of genes that are highly penetrant for ASD suggests that MIA might also exert important influence on downstream transcriptional programs of such genes. Here we tested this hypothesis with two such genes, *FMR1* and *CHD8*^54, 55^, because both are highly penetrant for ASD and are key master regulators of important neurodevelopmental processes including mRNA translation, transport, or localization (*FMR1*)^56, 57^ and chromatin remodeling (*CHD8*) of hundreds of genes implicated in transcription, cell division, proteolysis, DNA integrity, and signal transduction^51^. Interestingly, both *FMR1* and *CHD8* themselves are not dysregulated by MIA. However, this allows for an interesting test of the hypothesis that although these key genes are not directly dysregulated by MIA, their downstream targets may still be impacted by MIA and show evidence of substantial enrichment.

Across two FMRP target sets^56,57^ (see Supp Table 1 for gene lists) we found that MIA-downregulated genes are highly enriched in FMRP targets (Darnell targets: OR = 7.81, p = 2.56e-127; Ascano targets: OR = 2.07, p = 1.75e-30). Of the MIA-upregulated genes, enrichment was apparent in one of the two FMRP target lists (Darnell targets: OR = 0.22, p = 1; Ascano targets: OR = 1.86, p = 1.15e-21) (Fig 2B). For CHD8 targets, we also examined two target sets derived from either midgestational human fetal brain tissue and human neural stem cells^58^ or from human neural progenitor cells^59^ (see Supp Table 1 for gene lists). The MIA-downregulated (OR = 2.10, p = 3.52e-25), but not MIA-upregulated (OR = 1.19, p = 0.84) genes were enriched in CHD8 targets identified in midgestation fetal brain tissue and human neural stem cells^58^. In human neural progenitor cells both MIA-downregulated (OR = 1.66, p = 1.58e-7) and upregulated genes (OR = 2.71, p = 8.29e-87) were enriched in CHD8 targets ^59^ (Fig 2B). Overall, this evidence supports our hypothesis that while MIA does not directly affect *FMR1* or *CHD8*, two key genes with important transcriptional regulatory effects, it does potentially disrupt the same pathways by hitting their downstream targets.

### MIA-dysregulated genes are also dysregulated in child and adult ASD cortical transcriptome

We next examined the hypothesis that MIA-dysregulated genes are also dysregulated in in the child and adult ASD cortical transcriptome. To examine this, we re-analyzed two prior post-mortem ASD datasets from Voineagu et al.,^46^ and Gupta et al.,^45^. We found that ASD-downregulated genes in both datasets are substantially enriched in MIA-downregulated genes (Voineagu OR = 1.21, p = 0.047; Gupta OR = 2.12, p = 4.36e-23; see Fig 1C-D and Supp Table 1 for gene lists). These commonly downregulated genes are significantly enriched in processes such as transport_synaptic vesicle exocytosis, development_neurogenesis_synaptogenesis, and cell adhesion_synaptic contact (Fig 1E-F). Similar to downregulated genes, ASD-upregulated genes in both datasets were significantly enriched in MIA-upregulated genes (Voineagu OR = 1.57, p = 2.84e-5; Gupta OR = 1.57, p = 9.62e-7; see Fig 2C-D and Supp Table 1 for gene lists). Genes commonly upregulated in MIA and ASD were enriched in translation initiation processes (Fig 2E-F). However, despite such statistically significant enrichment, it is very noteworthy that a large majority of down- and upregulated genes perturbing prenatal developmental processes in MIA were not commonly dysregulated in the older child and adult ASD brain (Fig 1C-D; Fig 2C-D). That is, 92-96% of MIA-downregulated genes and 94-96% of MIA-upregulated genes were not commonly dysregulated in older child and adult ASD cortical tissue. Thus, while a specific subset of genes are commonly dysregulated by MIA in early fetal development and in older children and adult ASD cortical tissue, many other MIA-dysregulated processes in fetal development are likely not captured in common by looking at older ASD cortical tissue far beyond critical prenatal stages of brain development (e.g., upregulated cell cycle processes with likely role in neurogenesis).

### Translation and synaptic gene co-expression networks are highly preserved across MIA and ASD

We next tested whether systems-level transcriptome disruptions in MIA and ASD cortex are significantly similar or ‘preserved’. This approach goes beyond identifying overlap at the level of single genes and provides information about larger systems-level organization of the transcriptome and whether such dysregulated organization is similar across MIA and ASD cortical transcriptomic datasets. To do this, we implemented weighted gene co-expression network analysis (WGCNA) to identify preservation of systems-level structure of gene networks in MIA and ASD cortical transcriptome datasets^60,61^. We specifically examined ASD co-expression modules for on-average differential expression (DE) in module eigengene (ME) variation (i.e. systematic up or downregulation along the main principal axis of variation for a given gene module) and determined whether such DE modules were preserved in network structure in MIA. Co-expression modules that are both dysregulated and highly preserved across both datasets are ideal candidates for pinpointing common systems level biological disruption in both ASD and MIA.

We identified four consensus modules in ASD, M25, M3, M9, and M13, that show replicable on-average differential expression in both the Voineagu and Gupta datasets and also showed moderate levels of preservation in the MIA dataset. M25 was replicably upregulated in post-mortem ASD cortical tissue and was heavily enriched in translation initiation and translation elongation-termination (Fig 3A). M25 was the top hit in terms of preservation median rank and was the most preserved of any of the replicable DE modules with Zsummary preservation statistics of 8.5 and 8.8 (indicating ‘moderate’ preservations) respectively across Voineagu and Gupta ASD datasets (Fig 3C). Modules M3, M9, and M13 were replicably downregulated in Voineagu and Gupta ASD datasets and were enriched in a variety of synaptic functions (Fig 3B). These modules also showed moderate levels of preservation primarily with the Zsummary statistics above 2 (Fig 3D). These results further strengthen the evidence that MIA dysregulates systems-level structure of transcriptome in a manner similar to the dysregulation present in the ASD cortical transcriptome, with emphasis on upregulation of translation initiation processes as the strongest preserved signal across MIA and ASD.

**Figure 3.**
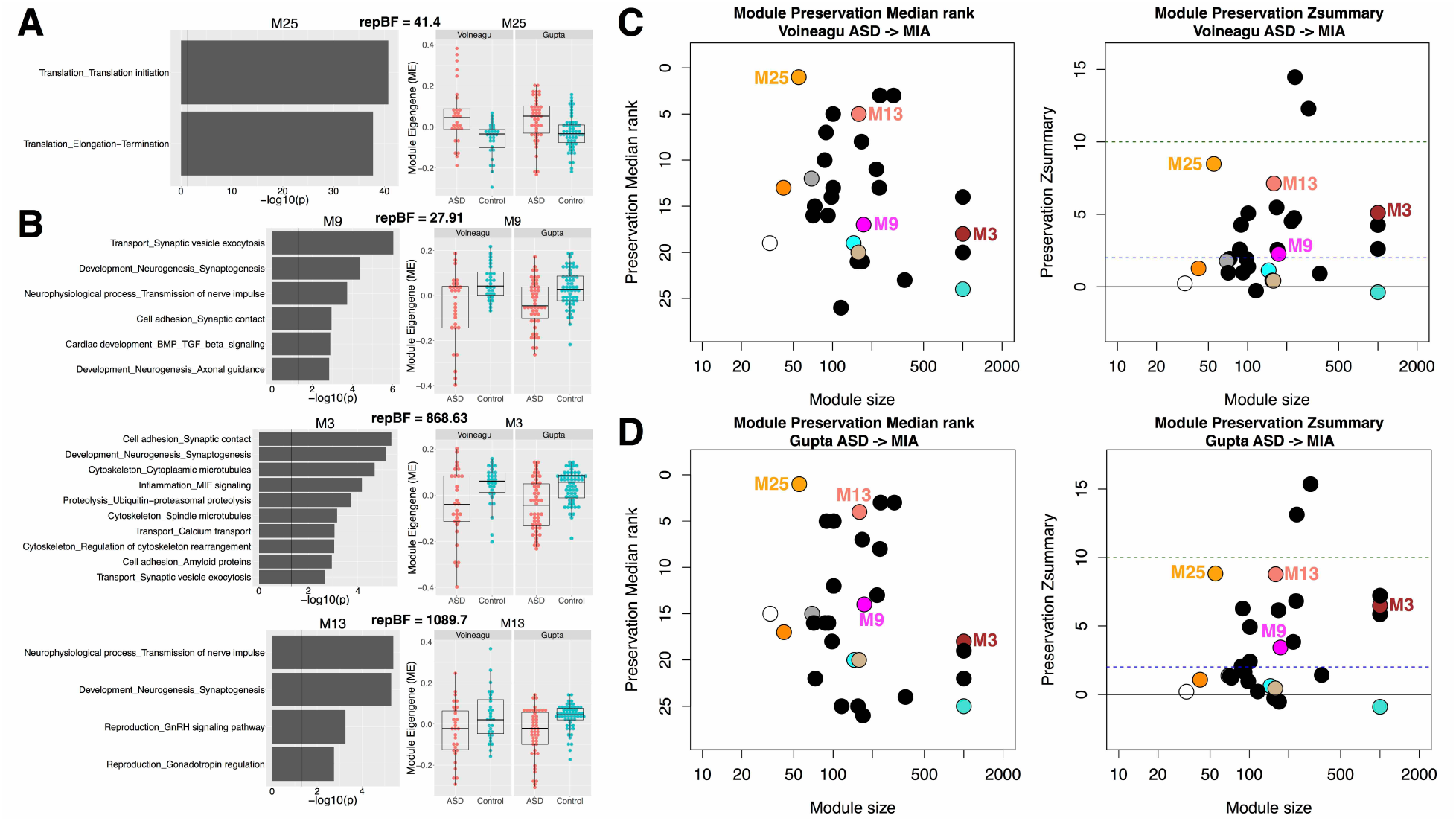
Preservation of dysregulated gene co-expression network organization across MIA and ASD. This figure shows ASD co-expression modules that are replicably dysregulated in ASD and preserved in network-level structure in the MIA dataset. Panels A and B depict gene co-expression modules that are replicably upregulated (A) or downregulated (B) in ASD cortical gene expression datasets. Scatter-boxplots show module eigengene (ME) expression levels with individual dots for each sample and boxplots that show the median and interquartile range (IQR; Q1 = 25^th^ percentile, Q3 = 75th percentile), as well as the outer fences (Q1 – (1.5*IQR) and Q3 + (1.5*IQR)). Next to each scatter-boxplot are results from process level enrichment analysis on each module. Above these plots are replication Bayes Factor statistics indicating evidence in favor of replication (repBF>10 indicates strong evidence in favor of replication). Panels C and D show module preservation statistics (median rank and Zsummary) for preservation between ASD cortical gene modules (C, Voineagu dataset; D, Gupta dataset) and MIA gene modules. The horizontal lines on the preservation Zsummary plot indicate categories for evidence of preservation, with Zsummary statistics between 2 and 10 indicating ‘moderate’ evidence for preservation. Modules represented by black dots are not differentially expressed between ASD and Control brains. Modules represented by colored dots (not black) and without a specific number (e.g. M25) are differentially expressed but not significantly preserved between ASD and MIA. Colored modules M25, M13, M3, and M9 are differentially expressed and significantly preserved between ASD and MIA.

### Activation of translation initiation processes dysregulates gene expression within members of the PI3K-TSC1/2-mTOR-EIF4E cascade in MIA and ASD

One common theme from the above differential expression and co-expression results of MIA and ASD transcriptomes is the presence of upregulated translation initiation processes. This common disruption suggests that either early environmental and/or genetic insults may lead to overlapping downstream effects via the dysregulation of translation pathways. Exaggerated cap-dependent translation is a well-known molecular mechanism regulating neurogenesis^62^ and contributing to synaptic and behavioral phenotypes associated with ASD and related neurodevelopmental disorders^63, 64^. Key to this mechanism is the aberrant regulation of the PI3KTSC1/2-mTOR signaling pathways, which in turn are responsible for the regulation of RPS6K1 and EIF4E-binding partners acting to promote translation initiation^65^. Enhanced mTOR signaling and cap-dependent translation initiation complex were found also in mouse models of human Fragile X syndrome characterized by the lack of FMRP^66-68^. Ultimately, these signaling pathways lead to the overexpression and activation of EIF4E-dependent mechanisms that have been directly linked to ASD both in mouse models and humans^69^. In support of this view, we have discovered evidence that downstream FMRP targets are dysregulated by MIA (Fig 2B) and that *EIF4E* displays the largest effect size of all ASD-associated genes (Cohen’s d = 8.27).

To further explore the effects of our network-based findings in MIA and its relevance to ASD, we asked whether the upregulated translation initiation-enriched ASD-module M25, which is the strongest preserved DE module in MIA, would affect members of the PI3K-TSC1/2-mTOR signaling pathways. To examine this hypothesis, we first constructed a network of M25 targets using the MetaCore canonical database and identified 3257 M25 direct targets. Importantly, functional analysis of these targets displayed a top enrichment in the regulation of cell cycle phase transition (G1-S and G2-M) and several developmental processes comprising several key regulatory genes (e.g. *AKT, JAK, NF-κB, PI3K, STATs, CDK, mTOR, NOTCH1, WNT and ERK/MAPKs*; Supp Table 2) as well as cap-dependent translation regulatory genes (*EIF4E* and its binding partners; Supp Table 2). These findings suggest that the ASD-upregulated translation initiation-enriched M25 module, which is preserved in MIA, may influence expression and activity of both PI3K-TSC1/2-mTOR signaling and EIF4E-dependent genes with predicted early neurodevelopmental effects on the timing of cell cycle phases during neural progenitor cell divisions.

To directly compare and quantify the effects of MIA and M25 dysregulation on this signaling pathway, we queried the MetaCore database to generate the shortest canonical network-path encompassing the PI3K-TSC1/2-mTOR-EIF4E axis. This network included 84 genes. We then asked the question of whether this axis is enriched in differentially expressed (DE) MIA genes, DE ASD genes, genes that are M25 targets, or ASD-associated genes (LGD, Missense, or SFARI) (see Figure 4). We found significant enrichments for genes DE in MIA (OR = 4.006, p = 7.14e-5) and for M25 targets (OR = 22.09, p = 1.89e-36) (Figure 4). Although only 14 of the 84 genes were DE in ASD brains resulting in a non-significant enrichment (OR = 1.47, p = 0.24), 10 genes of the 84 genes were those that are known to be ASD-associated (OR = 9.22, p = 4.62e-9) and of those 10, 7 were DE in MIA and/or ASD brains. Of note, we found that all key members of the signaling pathway (PI3K, TSC1, TSC2, mTOR, EIF4E) were dysregulated in MIA together with mTOR and EIF4E-binding partners (Fig 4). Altogether these findings provide compelling evidence that both MIA and ASD cortical transcriptome dysregulation involve the canonical PI3K-TSC1/2-mTOR axis regulating EIF4E-mediated cap-dependent translation.

**Figure 4.**
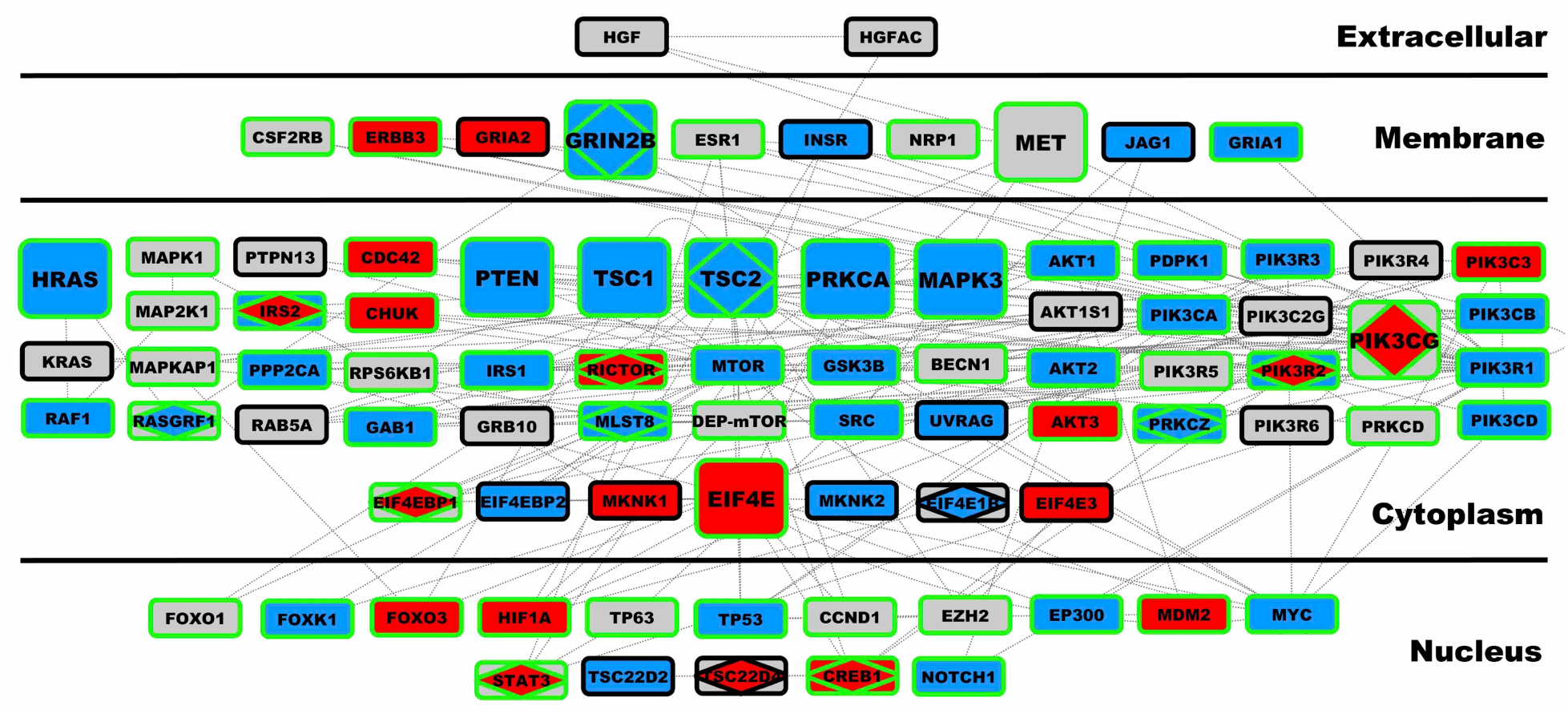
Canonical network encompassing the PI3K-TSC1/2-mTOR-EIF4E axis. This figure depicts all 84 genes comprising the shortest canonical network-path of the PI3K-TSC1/2-mTOR-EIF4E axis, as defined by the MetaCore GeneGO database. The plot is arranged by cellular compartment for each protein in the network. Nodes were depicted in larger size if they are members of the ASD-associated gene list compiled by SFARI Gene. Nodes with green borders are direct targets of the ASD co-expression module M25, which is ASD-upregulated and enriched in translation initiation and is preserved within the MIA dataset. Each node is colored on the inside to indicate directionality of differential expression (blue = downregulated, red = upregulated, grey = not differentially expressed). Rectangular shapes characterize all genes within this network. However, within each node a diamond shape indicate that the gene was differentially expressed in ASD brains.

### Cross-species validation of MIA-dysregulation of the Tsc1/2-mTOR-Eif4ebp1/2 axis

The network-level analyses suggest that MIA dysregulates the TSC1/2-mTOR-EIF4E signaling pathway. To directly test this hypothesis and validate inferences across model species, we performed a MIA mouse model validation experiment. Based on our findings in the rat model as well as literature evidence, we focused specifically on validating mRNA expression of the following targets: *Tsc1/2, mTor, Rps6k, Eif4e* and *Ei4ebp1/2* (EIF4E-binding proteins). Similar to the experimental design of the original rat MIA model dataset^26^, we induced MIA in pregnant dams using LPS at gestational day 12.5 in mice as previously described^70^ (see Methods). Mouse fetal brains were collected 2 hours post-LPS injection and mRNA transcript levels were quantified by qRT-PCR and expression levels were normalized by saline (SAL) controls. Consistent with the MIA rat gene expression findings, we replicated the effect of significant MIA-downregulation of *Tsc2* (*t* = −2.91, *p* = 0.012), *mTor* (*t* = −2.83, *p* = 0.012) *, Eif4ebp1* (*t* = −3.77, *p* = 0.0024) and *Eif4ebp2* (*t* = −5.05, *p* = 0.00078). We also replicated the MIA-induced upregulation of *Eif4e* (*t* = 2.13, *p* = 0.029). Replication of *Tsc1* MIA-induced downregulation was observed, albeit at trend-level significance (*t* = −1.70, *p* = 0.06), whereas *Rps6ka6* was not differentially expressed (*t* = −0.12, *p* = 0.54) (Fig 5). Together with the original discoveries in the rat MIA dataset, this cross-species validation strongly supports that MIA-induced transcriptional dysregulation of genes involves translation processes supported by the *Tsc1/2-mTor-Eif4ebp1/2* axis.

**Figure 5.**
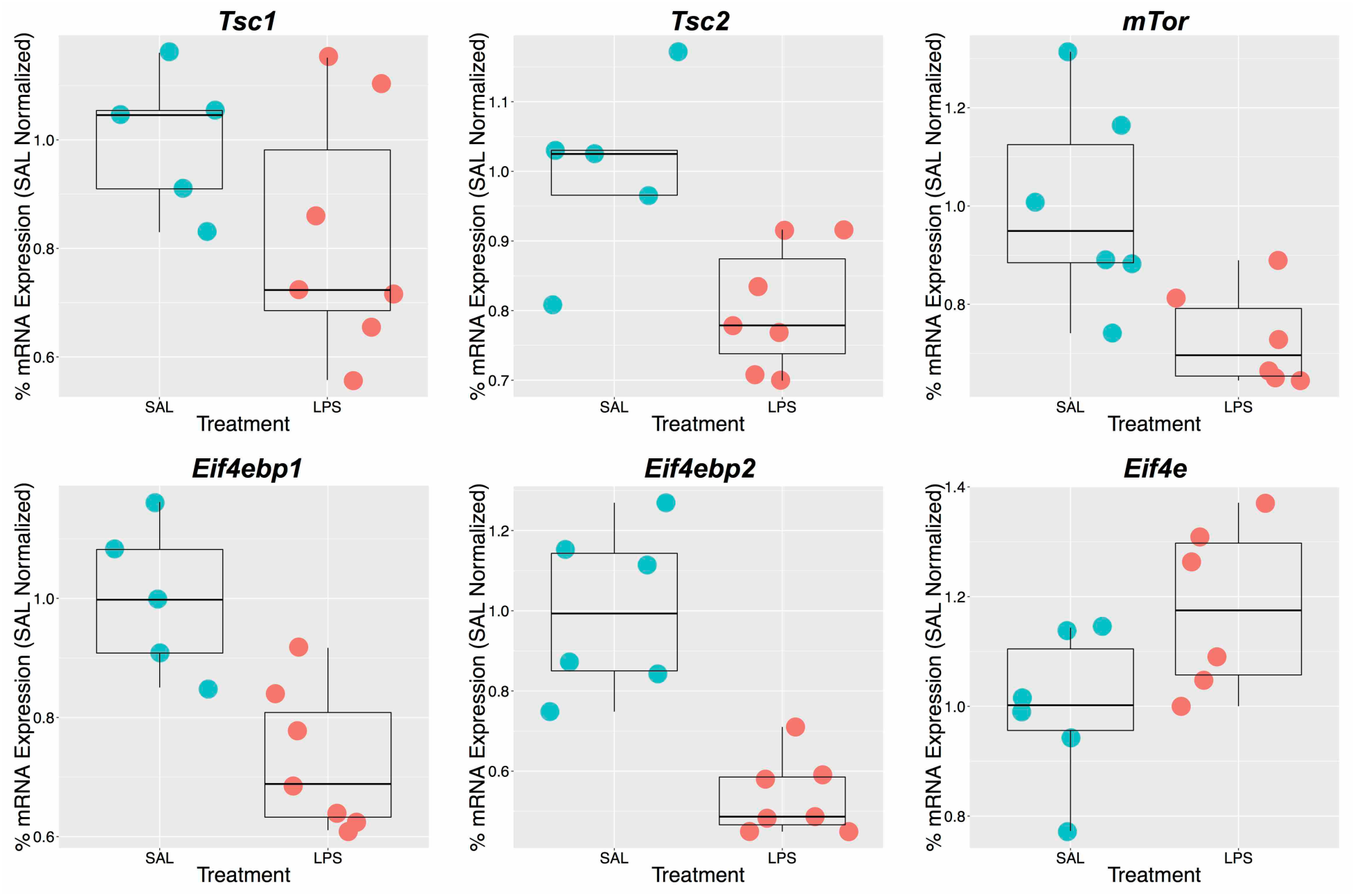
Cross-species validation of dysregulated gene expression within the Tsc1/2-mTor-Eif4ebp1/2 axis. This figure summarizes the results from qRT-PCR analysis of gene expression along the Tsc1/2-mTor-Eif4ebp1/2 axis hypothesized to show dysregulation from the previous rat model data. Each gene is displayed with a scatter-boxplot and gene expression for each individual sample is plotted on the y-axis as % mRNA expression normalized to the average value in the SAL condition. Boxplots show the median and interquartile range (IQR; Q1 = 25^th^ percentile, Q3 = 75^th^ percentile), as well as the outer fences (Q1 – (1.5*IQR) and Q3 + (1.5*IQR)).

## Discussion

In this study we examined animal models of MIA corresponding to late first trimester of gestation in humans, in order to better understand how MIA may lead to increased risk for ASD via impairments of early fetal brain development that are relevant to the pathophysiology behind ASD. We discovered that MIA induces dysregulation of the fetal brain transcriptome in several important ways. MIA downregulates expression of many genes known to be highly penetrant for ASD. Genes with smaller effect size for ASD risk were also downregulated by MIA though to a lesser extent than larger effect size genes. The strength of MIA-enrichment tended to follow the strength of evidence for association with ASD. This evidence suggests that MIA may have particular detrimental fetal programming impact for enhancing later risk for ASD by perturbing genes in early periods of fetal development that are of medium-to-high penetrance. MIA also significantly influences transcriptional programs that are downstream to highly penetrant mutations such as *FMR1* and *CHD8* genes, even when expression of these genes themselves may not be MIA-dysregulated. This evidence provides additional proof-of-concept that MIA-induced effects may converge to many of the same or overlapping pathways hit by some highly penetrant ASD mutations and this can occur without actual dysregulation of the target genes themselves. These findings bolster the intriguing possibility that MIA acts as an environmental etiological factor that disrupts specific key early developmental genomic mechanisms that are risks for ASD. MIA-induced disruption may work in a manner similar in directionality to rare highly penetrant ASD mutations. However, MIA may be different from such mutations in being a temporally transient event since it is restricted to circumscribed windows in fetal development rather than being persistent in disrupting protein synthesis over the entire lifespan as a deleterious mutation would be^71^. Nonetheless, in terms of the sheer number of genes affected by MIA, it could also be that such events may be more potent and common than germline mutations rarely found in ASD individuals. These results would further predict that when such environmental and gene mutation defects co-occur, aberrant effects could potentially be amplified towards one type of pathology or lead to more complex heterogeneous phenotypes. Prior work provides some evidence in support of such MIA-gene interactions, such as cooccurrence of MIA with *TSC2* haploinsuffiency^72^, rare de novo CNVs^73^, and *PTEN* mutations^34^. Intriguingly, both *TSC2* and *PTEN* are among the genes that are MIA-downregulated within the current dataset, which could increase abnormal neurogenesis. Overall, this work would suggest that MIA itself could constitute a sufficient environmental route through which the transcriptome in fetal brain development could be altered in ways similar to genetic etiologies associated with neural and behavioral phenotypes of ASD.

We also specifically examined how prenatal MIA-induced transcriptome dysregulation is similar to dysregulation of the ASD cortical transcriptome seen in later life. We found commonality between MIA and ASD in downregulation of synaptic-related processes and upregulation of translation-related processes. Interestingly, many of the genes and enrichment terms we found commonly downregulated are relevant to developmental processes, such as synaptogenesis, that occur at later prenatal and postnatal stages well after the point of the MIA event (i.e. see Supp Fig 1). One possible interpretation of this counterintuitive result could be that these downregulated genes play other important roles at earlier periods of first trimester brain development and that such roles are not well represented in current gene ontology annotations, especially when compared to their much more well-known canonical roles associated with synaptic processes in later development. This apparent pleiotropy of roles for these ASD-relevant genes is certainly an under-investigated area. For example, although many high-risk genes associated with ASD are commonly interpreted as being involved in later occurring processes such as neurite and synapse development, some research supports the idea that many of these genes also show prominent involvement in very early stages of brain development such as neural induction and early maturation of the neuroblast^74^. Thus, our results suggest an interesting new direction for future work that examines much earlier roles for ASD-associated genes, roles beyond those in later synaptic processes. Also important is that our results also show that the vast majority (approximately 92-96%) of MIA down- and upregulated genes that govern numerous key very early neural development processes (e.g., cell number, cell-type and laminar fate, migration, cell growth and differentiation) are *not* detected by examining differential expression of genes in cortical tissue in older child and adult ASD individuals. Therefore, it is important to consider that gene expression studies of the mature ASD cortex might, in fact, provide somewhat limited insight into the prenatal functional genomic pathology that may underlie the beginnings of ASD. As such, gene expression data from the mature ASD brain could be prone to false negatives and, as such, should be interpreted with caution. Indeed, a previous study showed age-dependent changes in abnormal cortical gene expression in ASD^47^.

A specific key dissimilarity between MIA in fetal development and the cortical transcriptome of ASD in children and adults is the presence of strongly upregulated cell cycle expression in MIA. This strong upregulation in fetal development is highly relevant given that the timing of MIA in this study occurs within a prominent time period for neurogenesis. In the child and adult ASD brain, there is a lack of any dysregulation in such cell cycle processes, but the developmental time period during which gene expression is assayed in later development corresponds to a period where neurogenesis processes are much less prominent. Thus, a potential major defect underlying early ASD development^13,31, 32, 51^ of dysregulated cell proliferation processes, likely cannot be adequately examined via study of the ASD brain in later development. We argue that MIA-upregulation of cell cycle processes and increased neurogenesis in early fetal development is likely a shared ASD-relevant aspect of pathophysiology^31,32^. Supporting this inference, we find that the ASD-upregulated M25 co-expression module that is preserved in MIA has strong downstream impact on cell cycle processes (see Supp Table 2) that are likely highly-relevant in early fetal development when neurogenesis is a highly prominent neurodevelopmental process.

We also found evidence for upregulation of translation processes in both MIA and ASD. Of note, transcriptional alteration of translation regulation was additionally supported by the FMRP target enrichment analysis that showed this is likely one of the strongest convergent signals in our comparative analysis of MIA and ASD effects on the cortical transcriptome. In prior work, we demonstrated upregulation of translation initiation in postnatal blood leukocyte expression in living ASD as compared to typical toddlers^75^. Furthermore, analysis of how gene co-expression modules interact within the cortical transcriptome of ASD (i.e. via eigengene network analysis of the same ASD transcriptome dataset analyzed here) supports the idea that a MIA-preserved translation initiation module is highly connected with immune/inflammation modules and other synaptic, cell cycle, and neurogenesis processes^76^. This systems biology link between atypical translational processes and neural dysregulation at the synapse, immune/inflammation processes and regulation of cell number is important to underscore, as it may suggest a larger more unified systems biological disruption than can be accounted for by only looking specifically at individual modules. In addition, the fact that dysregulation of translation initiation can be found systemically in blood^75^ and is not specific to neural tissue may allow for further investigation and hypotheses about the relation between this type of dysregulation with other sorts of systemic dysregulation and interaction with immune and inflammation processes.

Translation and protein synthesis mechanisms have been highly important within examination of syndromic forms of ASD. Kelleher and Bear suggested a ‘troubled translation’ hypothesis of ASD by linking mutations associated with syndromic forms of ASD to altered translation and disturbance of synaptic processes^63^. This hypothesis has been further elaborated by Santini and Klann and others, with new evidence supporting the crucial role of cap-dependent translation protein EIF4E^64,69, 77, 78^ in ASD pathophysiology^62^. The current data support these ideas that translation processes are integral to ASD and that MIA induces substantial early dysregulation of such processes. To investigate possible expression consequences of the upregulation translation processes seen in our analyses of MIA and ASD cortical transcriptomes, in our MIA mouse model experiment we tested whether key regulatory genes such as *TSC1/2* and *mTOR* that are upstream to the EIF4E-complex regulating cap-dependent translation, were a reproducible transcriptional phenotype of MIA. We report the novel finding that MIA in rodents during early fetal brain development influences the expression of the TSC-mTOR-EIF4E axis and the regulation of EIF4E binding proteins. This experimental evidence in a model system plus across species comparison of expression data indicates disrupted cap-dependent translation is common across rat MIA, mouse MIA and human ASD cortex. This finding warrants future examination to identify the specific downstream cellular and molecular phenotypes of cortical maldevelopment.

With regard to MIA-dysregulation of the TSC1/2-mTOR-EIF4E axis, we predicted that there are potential downstream effects involving genes that regulate cell proliferation, specifically controlling G1-S and G2-M cell cycle phases transition. The link we found between disrupted translation upon MIA or in ASD cortex and cell cycle processes is supported by other studies. MIA alters proliferation of cortical neural progenitor cells, laminar allocation of neurons, increased cortical thickness, increased cell density and patches of cortical dysplasia ^37 27, 33^. Increased proliferation of neural progenitor cells associated with brain overgrowth was observed after low-dose LPS treatment and was more pronounced in a *Pten* haploinsufficient background demonstrating clear genetic-environmental effects on early brain growth^34^. We recently demonstrated that genes frequently found mutated in ASD^1^ may regulate the downstream expression of genes directly relevant to brain size as well as other regulatory genes with cell cycle functions, particularly those involved in the regulation of the G1-S phase transition^51^. Further strengthening this evidence, *in-vitro* iPSC studies have bridged molecular and cellular phenotypes of cell cycle timing during neural progenitor cell division to abnormal cortical development in ASD subjects with enlarged brains^79,80^. Lastly, our evidence showing reproducible MIA-induced upregulation of EIF4E and downregulation of two binding proteins (EIF4EBP1/2) suggest a possible imbalance in the regulation of neurogenic versus radial progenitor divisions during development. In vivo evidence has indeed shown that normal expression and proper binding to EIF4E is required to maintain the correct balance^62^. We hypothesize that reduced binding of EIF4E partners may lead, directly or through a compensatory mechanism, to increase production of EIF4E which in turns is sufficient to abnormally expand the number of radial precursor cells^62^.

In summary, we show that many genes that are strongly dysregulated in early fetal brain development by MIA highly overlap with known ASD-associated genes and gene targets of two key ASD genes, *FMR1* and *CHD8*. At the same time, we show MIA additionally dysregulates large numbers of other genes that impact a multitude of early pivotal fetal programs that govern cell number, type, migration, laminar organization, axon guidance, growth and differentiation and these early functional genomic aberrances are largely *not* detectable at later ages in the mature ASD cortex. Increased awareness and knowledge about the impact of maternal infections during pregnancy on later risk for neuropsychiatric disorders like autism are particularly important given that such events are potentially preventable or could be largely reduced by changing practices^24,25^. In addition, MIA represents a potential etiology that could be more amenable to novel treatments^48, 49^. Our work here has highlighted the particular pathways related to translation initiation that could help to potentially explain the links between MIA and ASD and more work is needed to explore dysregulation of these processes and how potentially one could intervene and reshape such dysregulation. Finally, this work explains why MIA is a prominent risk factor for ASD and suggests that interactions between such risk and gene risk factors may enhance ASD risk.

## Methods

### ASD and MIA Cortical Transcriptome Datasets

The primary MIA dataset was a rat model microarray dataset downloaded from Gene Expression Omnibus (GEO; Accession ID: GSE34058) and was previously published on by Oskvig and colleagues^26^. This dataset applied a lipopolysaccharide (LPS) manipulation for the MIA-inducing event on gestational day 15, which in humans corresponds to the near the end of the first trimester of pregnancy. Gene expression was measured at 4 hours post-LPS injection on Affymetrix Rat GeneChip^®^ 1.0 ST chips. Data were preprocessed from the raw CEL files with background adjustment, quantile normalization, and summarization of probe intensities on log2 scale, using functions from the MATLAB Bioinformatics toolbox (i.e. rmabackadj.m, quantilenorm.m, rmasummary.m). We also analyzed two ASD cortical transcriptome datasets. The first was a microarray dataset from Voineagu and colleagues^46^ (GEO Accession ID: GSE28521) comprising frontal (BA9) and superior temporal cortex (BA41/42) tissue. The second dataset was an RNAseq dataset from Gupta and colleagues^45^ comprising frontal (BA44; BA10) and occipital cortex (BA19) tissue (http://www.arkinglab.org/resources/). For each ASD dataset we utilized the already pre-processed and quality controlled datasets publicly available in order to be as congruent as possible with prior published work.

### Differential Expression (DE) Analyses

Differential expression (DE) analyses were performed in R. For the MIA rat dataset, we used sva^81,82^ and limma packages^83^ for the DE analyses. Specifically, we utilized sva to determine a number of surrogate variables for inclusion as covariates in linear models via limma. For the ASD datasets, we utilized linear mixed-effect models (i.e. lme function within the nlme R package) to model fixed-effect variables of diagnosis, RIN, age, sex, PMI and median 5-prime to 3-prime bias (specific to the Gupta dataset) as well as model the random-effect of brain region. False discovery rate (FDR) correction for multiple comparisons was achieved using Storey’s method for FDR control^84,85^ implemented by the qvalue function in R. The FDR q-threshold for the MIA dataset was set conservatively at q<0.01 and for the ASD cortical transcriptome datasets was set to q<0.05.

### Weighted Gene Co-Expression Network Analysis (WGCNA)

Analyses of gene networks organized by co-expression patterns was implemented with the WGCNA package in R^60^. For datasets with multiple probes per gene, we collapsed genes with multiple to one unique probe per gene by selecting the probe with the highest mean expression value across the full dataset as implemented with the collapseRows function in R^86^. For the MIA dataset, we ran a signed WGCNA analysis where the soft power threshold was set to maximize R^2^ scale-free topology model fit as it plateaued above 0.8 and thus was set to 22. Soft power thresholded adjacency matrices were then converted into a topological overlap matrix (TOM) and a TOM dissimilarity matrix (i.e. 1-TOM). The TOM dissimilarity matrix was then input into agglomerative hierarchical clustering using the average linkage method. Gene modules were defined from the resulting clustering tree and branches were cut using a hybrid dynamic tree cutting algorithm (deepSplit = 2)^87^. Modules were merged at a cut height of 0.2 and the minimum module size was set to 30. For each gene module a summary measure called the module eigengene (ME) was computed as the first principal component of the scaled (standardized) module expression profiles. Genes that cannot be clustered into any specific module are left within the M0 module, and this module is not considered in any further analyses. For the ASD datasets, we ran a signed consensus WGCNA analysis in order to detect consensus modules for cross-dataset comparisons (implemented with the blockwiseConsensusModules function)^88^. All of the parameters were set identically to the MIA analysis except for the soft power thresholds, which were set to 14 for both datasets, based on similar criteria of maximizing R^2^ scale-free topology model fit. To test for differential expression at the level of ME variation we used linear mixed-effect models identical to those implemented in the DE analyses (i.e. same fixed and random effects). To identify MEs with replicable differential expression across both ASD datasets, we utilized t-statistics from the linear mixed models to compute replication Bayes Factor (repBF) statistics^89^ that quantify evidence for or against replication (see here for R code: http://bit.ly/1GHiPRe). Replication Bayes Factors greater than 10 are generally considered as strong evidence for replication. To identify replicable modules we first considered modules that possessed a significant effect passing FDR^84^ q<0.05 within the Voineagu dataset and then also required these modules possess significant effects in the Gupta dataset (FDR q<0.05) and that this evidence quantitatively produces evidence for replication with a replication Bayes Factor statistic > 10. To test ASD gene modules for preservation with the MIA dataset we ran a module preservation analysis using the function modulePreservation and set the number of permutations to 200.

### MetaCore GeneGO Enrichment Analyses

In order to understand what molecular processes our gene lists were enriched in, we used MetaCore GeneGO software platform (https://portal.genego.com/) to perform all enrichment tests. These analyses were done at the level of ‘Process Networks’ within MetaCore.

### Gene Set Enrichment Analyses

All gene set overlap analyses were implemented using the sum(dhyper()) function in R. The background set size for all enrichment analyses was set to the total number of probes within the MIA dataset (i.e. 22,071).

### Network Analysis of PI3K-TSC1/2-mTOR-EIF4E Axis

The analysis of the predicted targets of module M25 dysregulation was performed by querying the Metacore GeneGO database (https://portal.genego.com/). All genes from the M25 module (61 genes) were used as bait to search the database for canonical interacting partners using one interaction distance (the “no filtering” option was used). This search yielded networks with 3257 genes in total. We saved this gene list and ran enrichment analysis to learn about the biological processes possibly affected by the M25 dysregulation.

Similarly, to quantify the predicted effects of MIA and M25 dysregulation on the PI3KTSC1/2-mTOR-EIF4E signaling pathway we queried the Metacore GeneGO database to identify the shortest canonical network connecting these five key regulatory genes. These genes were thus used as bait and the shortest canonical network with a maximum number of 2 steps in the path was selected. Eighty-four nodes and their interactions were exported from Metacore and the canonical network was reproduced in Cytoscape (http://www.cytoscape.org). Exporting the network in Cytoscape facilitated the color-coding of the genes to display the overlap with the M25 targets and the differentially expressed genes from the MIA and ASD cortices as well as the ASD-associated genes.

### Mouse MIA Model Experiment

All animal studies were performed in accordance with NIH guidelines for the use of animals and all procedures were reviewed and approved by the Stanford Institutional Animal Care and Use Committee. Timed pregnancies of C57BL/6J mice were obtained by housing a female and a male overnight. The individual mouse was separated the next morning and defined the mid-day of that day as embryonic day 0 (E0.5). The pregnant females were identified by body weight gain during the time course of pregnancy. To induce MIA responses, at E12.5, the pregnant dams were injected intraperitoneally with lipopolysaccharide (LPS) from *Escherichia coli* 055:B5 (L4524, Sigma-Aldrich, St. Louis, MO) at doses 60 µg/kg dam’s body weight. Control dams were injected with saline (SAL, vehicle) only.

### qRT-PCR Analysis

Whole fetal brains were homogenized in TRIzol reagent (Life Technologies, Carlsbad, CA) using RNase-free disposable pestles (Kimble Chase, Vineland, NJ) to extract total RNA. Following chloroform, 100% ethanol was added to precipitate the aqueous phase containing RNA. Then the aqueous phase was transferred onto a QIAgen RNeasy mini spin column and RNA/DNA was isolated with the QIAgen RNeasy mini kit (QIAGEN, Venlo, Netherlands) following the manufacturer’s protocol. DNA was digested using DNase-I enzyme (QIAGEN) for 15 min at RT. Nanodrop spectrophotometer (ThermoFisher Scientific, Waltham, MA) was used to assess the quality and concentration of isolated RNA. RNA was stored at −80°C until cDNA synthesis. To synthesize cDNA template, reverse transcription PCR reaction was performed on extracted RNA with MultiScribe reverse transcriptase (ThermoFisher Scientific, Waltham, MA) and random primer sets in the following condition; 25°C 10 min, 37°C 120 min, 85°C 5 min. The synthesized cDNA was kept at −20°C until qRT-PCR reaction. VeriQuest Probe qPCR Master Mix (Affymetrix, Santa Clara, CA) was used in the following qPCR reaction with Fast Real-Time PCR systems (Applied Biosystems, AP7900HT, Waltham, MA); 50°C 2 min, 95°C 10 min, 40 cycles of (95°C 15 sec, 60°C 1 min). The FAM-conjugated TaqMan qRT-PCR primer sets used in the present study and ROX was used as reference. Percentage of mRNA expression was calculated by converting relative mRNA copy number from differences between Ct (Cycle threshold) values of *Gapdh* (a housekeeping gene) and the gene-of-interest. The relative mRNA expression levels in LPS-treated fetal brains was normalized by SAL-treated control levels.

#### Taqman qRT-PCR Primers

| Gene | Ref Seq number | Catalog number | Exon Boundary |
| --- | --- | --- | --- |
| *Gapdh* | NM_001276655.1 | Mm99999915_g1 | <u>2-3</u> |
| *Tsc1* | NM_001289575.1 | Mm00452208_m1 | <u>3-4</u> |
| *Tsc2* | NM_001039363.2 | Mm00442004_m1 | <u>16-17</u> |
| *Eif4e* | NM_007917.3 | Mm01621873_s1 | <u>8-8</u> |
| *Eif4ebp1* | NM_007918.3 | Mm04207378_g1 | <u>2-3</u> |
| *Eif4ebp2* | NM_010124.2 | Mm01149891_m1 | <u>1-2</u> |
| *mTor* | NM_020009.2 | Mm00444968_m1 | <u>6-7</u> |
| *Rps6ka6* | NM_025949.3 | Mm01225184_m1 | <u>22-23</u> |

All statistical tests on qRT-PCR data employed one-tailed independent samples *t*-tests that do not assume equal variances (i.e. t.test function in R). The one-tailed predictions are justified by the directionality of differential expression observed in the rat microarray MIA dataset. Control for multiple comparisons was achieved by setting the FDR threshold to q<0.05.

## Acknowledgments

We would like to thank Nathan Lewis and Anthony Wynshaw-Boris for helpful discussion of this work. This work is supported by grants (KL2TR00099 and 1KL2TR001444) from the University of California, San Diego Clinical and Translational Research Institute to Dr. Pramparo and a grant from the Simons Foundation Autism Research Initiative awarded to Prof. Courchesne (SFARI #176540) and Prof. Theo D. Palmer (SFARI #323220), and Postdoctoral fellowship from Child Health Research Institute at Stanford University School of Medicine to Dr. Moon.

## Author Contributions

MVL, EC, and TP conceived the idea for the study. MVL and TP conceived and implemented all data analyses. HMM, JS, TDP, and TP conceived the idea for MIA mouse model validation study and performed qRT-PCR experiment/data analysis. MVL, EC, and TP interpreted the results and wrote the manuscript. All authors read and approved the final manuscript.

## Supplementary Information

**Supplementary Figure 1: Schematic timeline of prominent processes occurring in fetal brain development**

This figure contains a schematic of prominent processes occurring during different periods of human fetal brain development. For this study, the MIA-inducing event occurs near the end of the first trimester.

**Supplementary Table 1: Gene lists**

Contains gene lists for differentially expressed genes in MIA and ASD datasets, ASD-associated gene lists for enrichment tests, and gene lists for coexpression modules that are dysregulated in ASD and preserved in MIA.

**Supplementary Table 2: Enrichment tables**

Contains enrichment results for MIA down- and upregulated genes and M25 targets.

